# Relationship between the Changes over Time of Bone and Muscle Health in Children and Adults: A Systematic Review and Meta-Analysis

**DOI:** 10.1101/541524

**Authors:** Locquet Médéa, Beaudart Charlotte, Durieux Nancy, Reginster Jean-Yves, Bruyère Olivier

## Abstract

Various cross-sectional studies provide an abundance of evidence that shows a relationship between bone quantity and muscle health. However, one question remains, less-often studied: is their development - or decline – associated? The aim of the research was to conduct a systematic review and meta-analysis to summarize the studies exploring the association between changes in bone mineral density (BMD) and changes in muscle parameters (registration CRD42018093813). We searched for prospective studies, both in children and adults, by consulting electronic databases (Ovid-MEDLINE, Ovid-AMED, Scopus). Each review steps were performed by two independent reviewers. For outcomes reported by less of 3 studies, we synthetized the results narratively. In other cases, a meta-analysis was performed, giving an overall r coefficient and its 95% confidence interval (CI). Fifteen papers were included. In connection with the change of BMD, 10 studies concerned the parallel change of lean mass, 4 were about grip strength, and 1 was about physical performance. Children were the population of interest for 5 studies, while the aging population was the focus of the other studies. The correlation between hip BMD and lean mass was significant, with an overall coefficient r = 0.37 (95% CI 0.23-0.49). High heterogeneity was observed between studies but the length of follow-up, sex and study quality did not seem to significantly influence results. The systematic review allowed some other highlights: a significant link between changes in BMD and changes in muscle strength was observed (p-value <0.05 in the 4 studies), in addition to changes in performance (1 study, r = 0.21, p-value = 0.004). Despite the heterogeneity between studies, we highlighted a significant association between the change of BMD and the change of various muscle parameters. Future studies should investigate preventive and therapeutic strategies that are based on a single entity: the ‘muscle-bone unit’.

## Introduction

A good musculoskeletal health is crucial throughout the course of life. In childhood, it allows an optimal development [1], since the acquisition of bone mass is significantly impacted by the muscle function [2]. In adulthood, good musculoskeletal health is necessary, especially to prevent disorders affecting quality of life [3]. Then, in the aging process, optimal musculoskeletal health prevents the loss of functional performance and subsequently allows for better autonomy and preserve independence [4]. Indeed, direct and harmful consequences of decreased musculoskeletal health exist, including sarcopenia and osteoporosis, resulting in an increased propensity for falls and fractures, hospitalizations and, ultimately, premature death [5–10].

A body of evidence proves that the two entities of bone and muscle are highly linked due to their many interactions and interconnections. Obviously, the most noticeable link is demonstrated by the mechanical influences between these two tissues in the theory that was developed by Harold Frost [11]: There is a mechanical loading that is generated by the movement of the muscle on the bone, leading to a positive relationship between the lean masses and bone tissues. Dealing with this ‘mechanostat’ principle and on the “bone-muscle” unit in children and adults, different modulators can be considered, such as hormonal, nutritional, behavioral or environmental factors [12,13].

From a clinical point of view, the link between lean mass and bone has already been demonstrated in many cross-sectional studies, as synthesized by a systematic review conducted in 2014 [14]. Indeed, this work concluded that lean mass was significantly associated with bone mineral density (BMD), estimating an overall correlation coefficient of 0.39 (95% confidence interval (CI) 0.34-0.43). With regard to muscle strength, its significant cross-sectional link with BMD has already been established a few times in postmenopausal women [5–7], but the research regarding childhood or adulthood is much less plentiful. There is a similar finding regarding the link between BMD and muscle performance: some studies, especially those of participants over the age of fifty, showed a significant but moderate correlation between the quantity of bone and the physical performance of subjects [8–10].

However, fewer studies have investigated the longitudinal evolution of BMD with its parallel evolution of lean mass, muscle strength and physical performance. Therefore, we systematically recorded longitudinal studies exploring the relationship between the changes of BMD and the changes in muscle components (i.e., lean mass, muscle strength and physical performance) with the ultimate goal to synthetize the findings of each independent study using a narrative review or a meta-analysis.

## Methods

Each stage of our systematic review and meta-analysis rigorously respected the guidelines proposed by the Preferred Reporting Items for Systematic Reviews and Meta-Analysis (PRISMA)[21]. Our issue of interest was first correctly identified and defined using the following PICOS strategy: Population or disease - bone mass; Intervention - the effect of the passing of time (growth in the child, advanced age in the adult and the elderly); Comparator - Not applicable; Outcomes - Lean mass, muscle strength and physical performance; Study design - Prospective. Our goal was therefore to systematically search and summarize the studies describing the association between the changes in bone mass with regards to the changes in muscle function (i.e., lean mass, muscle strength and physical performance). A protocol has been developed and published on PROSPERO under the registration number CRD42018093813. Subsequently, we went through the different stages detailed below: literature search, study selection, data extraction, quality evaluation, data synthesis and statistical analysis.

### Literature Search Strategy

The Ovid-MEDLINE (1946 to August 2017), Scopus (1960 to August 2017) and Ovid-Allied and Complementary Medicine Database (AMED) (1995 to August 2017) electronic databases were searched in August 2017, with an update in December 2018, to identify the relevant studies that assessed the prospective association between changes in bone mass and in muscle components. Additionally, a search of systematic reviews and other syntheses of previous literature was also initiated to manually identify potentially relevant scientific references in the bibliography using the Ovid-Cochrane Database of Systematic Reviews, Ovid-ACP Journal Club and Ovid-DARE databases. The main keywords that were employed comprised terms as ‘Bone Mineral Density, ‘Lean mass’, ‘Muscle Strength’, ‘Physical Performance’, and ‘Prospective Study’. No limitation for the date was applied, but a restriction was set for English or French languages. The detailed search strategy with the key words that were applied in the Ovid interface is available in Table 1.

**Table 1.** Search Strategy applied via Ovid

|  |  |  |  |
| --- | --- | --- | --- |
| 1 | Bone Density/ | 29 | (Musc* adj2 Disabilit*).ti,ab. |
| 2 | (Bone adj2 Densit*).ti,ab. | 30 | Sarcopenia/ |
| 3 | (Bone adj2 Mineral adj2 (Densit* or Content*).ti,ab. | 31 | Sarcopenia.ti,ab. |
| 4 | Osteoporosis/ | 32 | exp Muscle Strength/ |
| 5 | Osteoporos*.ti,ab. | 33 | (Muscle* adj2 Strength*).ti,ab. |
| 6 | Osteoporosis, Postmenopausal/ | 34 | (Hand adj2 Strength*).ti,ab. |
| 7 | Osteoporos* Postmenopausal.ti,ab. | 35 | (Grasp* adj2 Strength*).ti,ab. |
| 8 | Bone Diseases, Metabolic/ | 36 | (Grip* adj2 Strength*).ti,ab. |
| 9 | (Bone adj2 Metabolic adj2 Disease*).ti,ab. | 37 | (Pinch* adj2 Strength*).ti,ab. |
| 10 | Osteopenia*.ti,ab. | 38 | (Physical adj2 Performance*).ti,ab. |
| 11 | Bone Demineralization, Pathologic/ | 39 | (Physical adj2 Endurance*).ti,ab. |
| 12 | Bone Demineralization*.ti,ab. | 40 | (Endurance* adj2 Physical adj2 Activit*).ti,ab. |
| 13 | (Bone adj2 Loss*).ti,ab. | 41 | (Physical adj2 Abilit*).ti,ab. |
| 14 | (Bone adj2 Decline*).ti,ab. | 42 | (Physical adj2 Function*).ti,ab. |
| 15 | (Bone adj2 Weakness*).ti,ab. | 43 | or/18-42 |
| 16 | (Bone adj2 Wasting).ti,ab. | 44 | Cohort Studies/ |
| 17 | or/1-16 | 45 | Cohort Stud*.ti,ab. |
| 18 | (Musc* adj2 Tissue*).ti,ab. | 46 | Cohort Analys*.ti,ab. |
| 19 | (Muscle adj2 Mass*).ti,ab. | 47 | Follow-Up Studies/ |
| 20 | (Lean adj2 Mass*).ti,ab. | 48 | Follow Up Studi*.ti,ab. |
| 21 | (Lean adj2 Body adj2 Mass*).ti,ab. | 49 | Longitudinal Studies/ |
| 22 | (Lean adj2 Tissue*).ti,ab. | 50 | Longitudinal Stud*.ti,ab. |
| 23 | (Fat Free adj2 Mass*).ti,ab. | 51 | Prospective Studies/ |
| 24 | (Fat Free adj2 Body adj2 Mass*).ti,ab. | 52 | Prospective Stud*.ti,ab. |
| 25 | (Muscle* adj2 Loss*).ti,ab. | 53 | Prospective Change*.ti,ab. |
| 26 | (Muscle* adj2 Decline*).ti,ab. | 54 | or/44-53 |
| 27 | (Muscle* adj2 Weakness*).ti,ab. | 55 | 17 and 43 |
| 28 | (Muscle* adj2 Wasting).ti,ab. | 56 | 54 and 55 |
|  |  | 57 | limit 56 to (english or french) |

### Study Selection

A first screening step, performed by two independent investigators, was based on title and abstract of each reference that was yielded by the literature search. This procedure allowed for the exclusion of irrelevant studies according to the strict eligibility criteria that are shown in Table 2. Mainly, inclusion criteria included: (1) Prospective studies, (2) interested about changes in bone mineral density and (3) changes in muscle function (i.e., lean mass, muscle strength or physical performance). For the second stage, the two investigators independently read the full texts of the articles that were selected by the initial screening, and they scrutinized the inclusion and exclusion criteria for the identified studies. Doubts and differences of opinion about a potential inclusion were settled following a discussion between the researchers, with the intervention of a third if necessary.

### Data Extraction

Relevant data were independently extracted by the two reviewers according to a standardized data extraction form, which was previously pretested on a sample of two studies. Discrepancies between the two collaborators were solved by discussion, with mediation from a third peer if needed. The following data were extracted: first author; journal name; year of publication; country; study objective; sociodemographic data; sample size; time of follow-up; tools and cut-offs used to assess bone and muscle components; statistical outcomes (i.e., correlation coefficient or β coefficient); adjustment factors; conclusion; potential conflicts of interest and funding. Authors and coauthors were systematically contacted by email if one of these data were unavailable in their report.

### Quality Assessment

All included studies were appraised for their methodological quality by two independent reviewers using the Newcastle-Ottawa Scale (NOS) for cohort studies; this scale was composed of three grades: group selection, comparability, and exposure and outcome assessment. A maximum of 9 points could be granted, with a score of 9 thus representing the highest methodological quality. For this evaluation, disagreements between the two reviewers were resolved thanks to the opinion and advice of a third expert.

### Data Synthesis and Statistical Analysis

A descriptive analysis of the included studies has been performed under the format of a narrative report. Some data that were required for the meta-analysis were missing. To counter this, first and/or last authors were contacted by email for additional information. For each association of changes in BMD and changes in lean mass that was reported by at least three papers depending on the site of measurement (i.e., total hip, femoral neck, lumbar spine or total body), a meta-analysis was undertaken, combining the statistical results of each study to determine an overall effect size, which was expressed as a correlation coefficient r with its 95% confidence interval (CI) and p-value, reported visually through a forest plot. A certain number of studies shared association results after their adjustments, meaning that these analyses computed in a multivariable model, yielding a β coefficient with its standard error. We have contacted all of the authors of the different studies that provided only the β coefficient to obtain the data concerning the coefficient of correlation r. Only one author sent us the new exact r value [22]. In the other cases, we computed the correlation coefficient r from the β coefficient, following the formula proposed by Peterson and Brown, in 2009 [23]. Indeed, as suggested by these authors, this method (i.e., finding the r from the β) generally generates an accurate estimate of the effect size, giving a definite advantage for completing the meta-analysis, considering that the sampling errors would be more numerous if we excluded studies simply because they did not report the correlation coefficient r [23]. Since we assumed a priori that the correlation estimates could fluctuate across studies because of a real association in each study but also by chance (and, subsequently, sampling variability), we used random-effects models to combine the results of the studies to find a pooled effect size. Moreover, the statistic test I², an estimator of inconsistency, as well as the χ² test allowed for us to explore the heterogeneity [24]. A one-way sensitivity analysis was also performed to evaluate the consistency of our results when omitting one study at a time, and we repeated this process for each study. A subgroup analysis was conducted because of the assumption of differences based on the life-course position of the individuals who were evaluated (i.e., children or older adults). Difference between groups was assessed by performing a Q-test based on analysis of variance. To investigate potential other sources of heterogeneity modifying the association between the evolution of bone mass in parallel with that of lean mass, we computed meta-regression models, which are composed of different moderators: mean age, quality of study, length of follow-up and sex of participants studied. To estimate the presence of a potential publication bias, we did not visually inspect the funnel plots since less than 10 studies were included in the different meta-analyses, but we instead used the Egger’s regression asymmetry test to detect it. In each statistical result, a p-value equal or less than the 5% critical level was considered as statistically significant. All processes that were undertaken for the meta-analysis were realized using the software package Comprehensive Meta-Analysis, version 2 (Biostat, USA).

## Results

### Rendering of our Literature Search

The search through the literature yielded 1889 relevant references, after removing 49 duplicates. We additionally found, by handsearch, 3 relevant studies. From these 1892 articles, after a long process of review and discussion (Figure 1), fifteen studies were finally included [11,22,25–37], detailed as follows:

- 10 studies examining the link between evolution of lean mass and evolution of bone mass;
- 4 studies focusing on the link between evolution of muscle strength and evolution of bone mass;
- 1 study dealing with the link between evolution of physical performance and evolution of bone mass.

**Figure 1.**
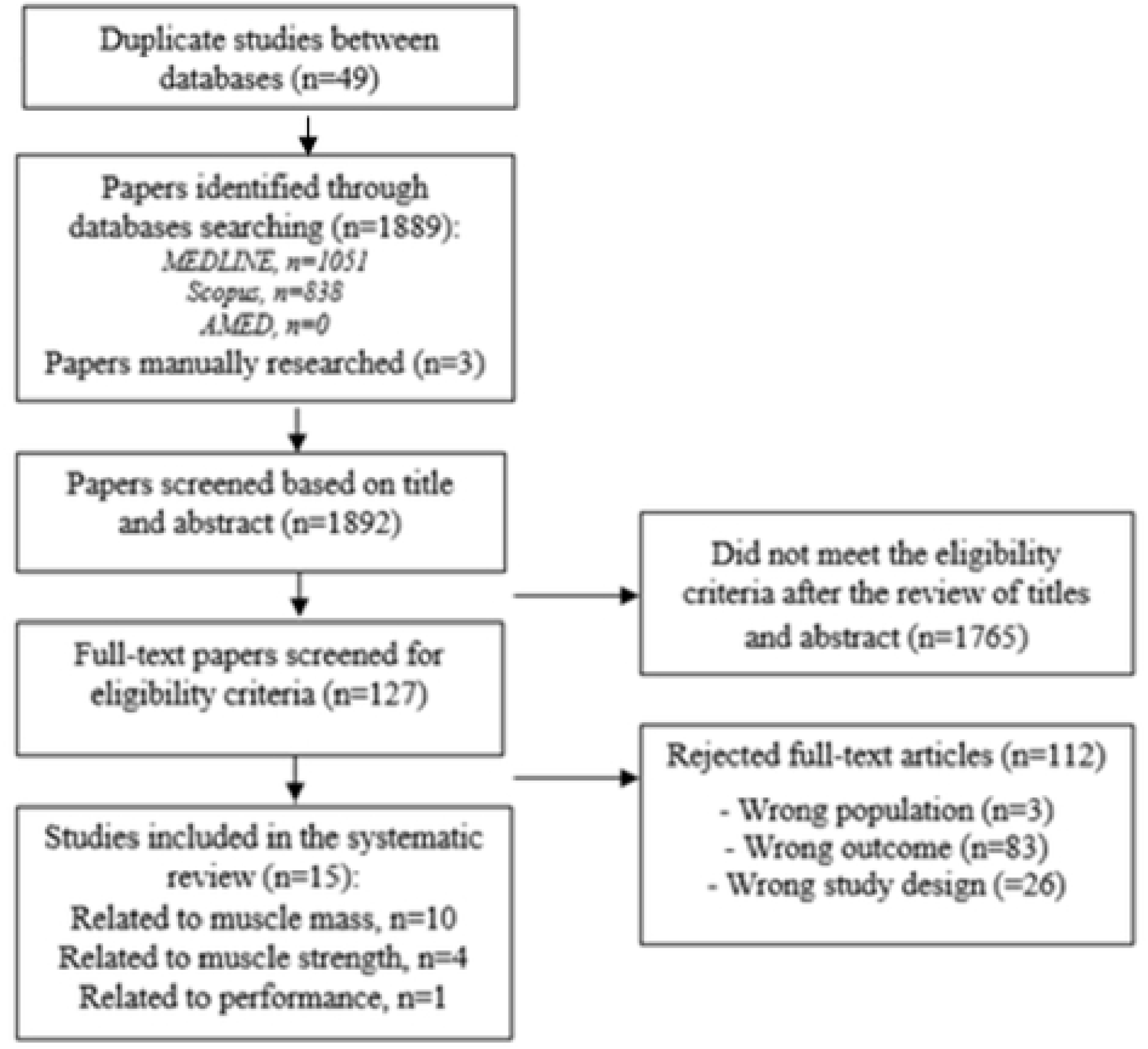
Detailed literature search flow diagram

At the end of December 2018, we made an update to our search and relaunched our search strategy within the databases. No new relevant references were identified for our subject of interest.

### Characteristics of the Included Studies

The general characteristics of the 15 references included are detailed in Table 3. These studies were published between 1997 and 2015. The population of interest was children for 5 studies [25–27,31,32], and the population of interest of the other studies were in female around the menopause age or the older age. With regards to the gender of the participants, 10 studies only focused on female subjects, 2 studies on male subjects only [27,38], and 3 other studies consisted of a population of both men and women [31,32,36]. The duration of follow-up of these longitudinal studies ranges from 1 year to 10 years. The instrument that was used to assess bone and lean mass was the Dual Energy X-ray Absorptiometry (DXA) device. Muscle strength was measured with a handgrip dynamometer [11,28,37] or a chair dynamometer [26] and the physical performance was established on the basis of a standard walk test of 5 meters [30]. One study presented conflicts of interest [22], 6 others recorded no conflict of interest, and 8 manuscripts did not mention the presence or absence of it. Additionally, 14 out of 15 studies are funded by a foundation, ministry, grant or national institute, and a single [39] study does not report whether or not they obtained a source of funding. The different studies are of variable quality (Table 4), from satisfactory to excellent.

### Relationship between Changes in Bone Mass and Changes in Lean mass

We have therefore identified 10 articles focusing on the parallel evolution of bone and lean mass. For the obvious reasons of clinical considerations, we separated our analysis in two ways:

- When the measurement of the BMD at different sites of the skeleton corresponding to the measurement of different bone compartments whose trabecular bone and cortical bone content was variable, we decided to perform a meta-analysis per measurement site of this BMD.
- The bone mineral content (BMC) can significantly differ from bone mineral density, so the study of Heidemann *et al*. (2015) [32], which focused only on BMC, was analyzed separately with a narrative report.

### Hip Bone Mineral Density and Lean Mass

The 8 analyzed studies [22,25–27,29,31,36,38] comprised a total of 2022 patients. Their quality varied by 5 points in the NOS score [25] under the maximum score of 9 points [38] (Table 4). Three studies reported an analysis of children’s health [25,27,31], and 5 studies were about postmenopausal or older subjects [22,29,36,38,39]. The association between the evolution of hip BMD and the evolution of lean mass was significantly correlated in all studies except for that of Chen *et al*. (1997) [39] (p-value >0.05) and Hrafnkelsson *et al*. (2015) [31] (p-value>0.05). The correlation coefficients r varied from 0.05 [39] to 0.69 [27] (Figure 2). Among these results, the correlation coefficients r were imputed from the available β coefficient for all of the studies except for that of Liu-Ambrose *et al*. (2006) [22] and that of Vicente-Rodriguez *et al*. (2005) [27]. A meta-analysis was performed to combine the results of these different studies (Figure 2).

**Figure 2.**
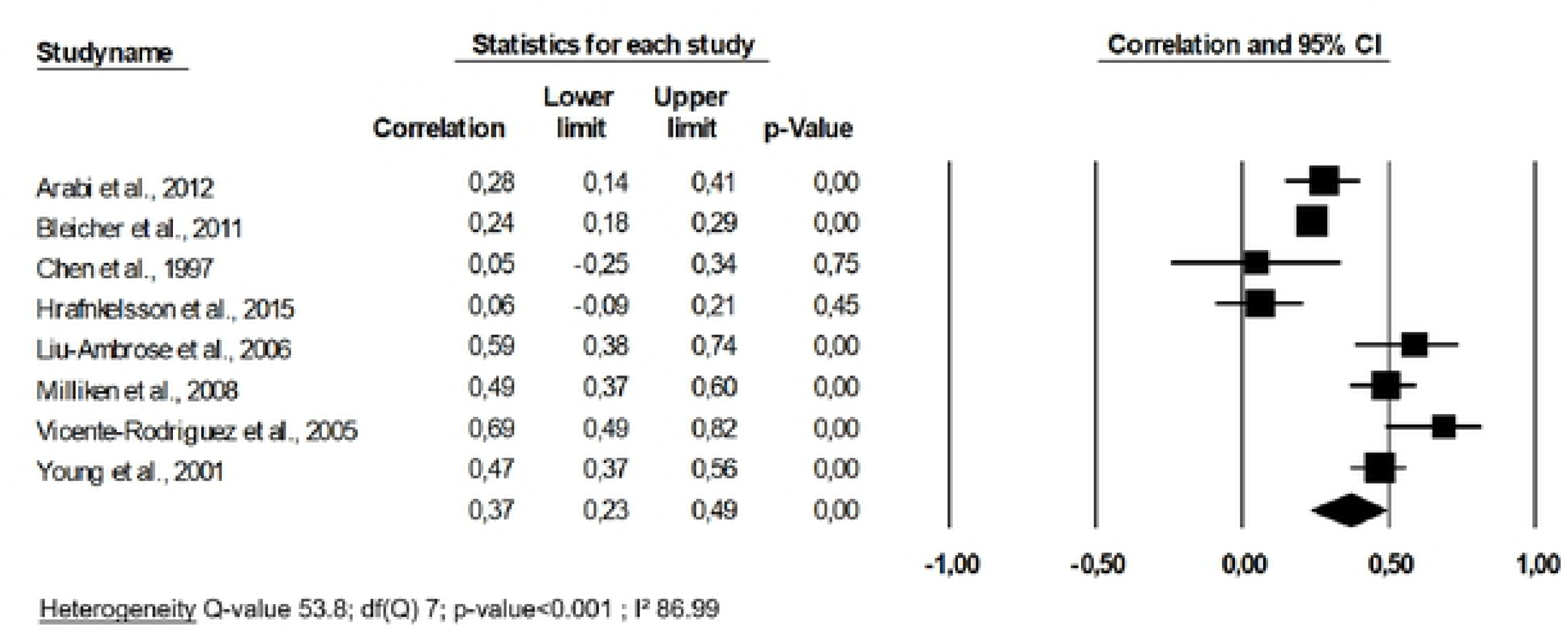
Association between changes in hip bone mineral density and changes in lean mass: a meta-analysis

An overall correlation coefficient r of 0.37 (95% CI 0.23-0.49, p-value <0.001) was yielded by pooling the results of the 8 studies, indicating that the prospective changes in hip BMD were significantly and moderately correlated to the changes in lean mass. The subgroup analysis indicated that there was a significant difference in effect size (p-value<0.001) between that of children (3 studies, r = 0.43 (95% CI: 0.06-0.69)) and that of adults and older individuals (5 studies, r = 0.34 (95% CI 0.19-0.48)). There was heterogeneity in our meta-analysis (I^2^ = 86.99, p-value<0.001), so a meta-regression was performed to explore this phenomenon. We found no significant impact at the level of the quality of study, the mean age analyzed, the sex or the duration of the follow-up on the overall effect size (all p-values>0.05). The Eggers’ regression analysis demonstrated that publication bias was not present (p-value=0.27). The one-way sensitivity analysis showed the consistency of our results, showing that they had roughly the same correlation coefficient, and the heterogeneity I^2^ value exceeded 85.75% in all cases.

### Femoral Neck Bone Mineral Density and Lean mass

Four studies of a total of 649 individuals investigated the relationship between femoral neck change and lean mass change [25,29,36,39], and the methodological quality ranged from satisfactory [25] to good [29,36,39]. The studies focused on adulthood (i.e., in this case, premenopausal women) except one that focused on childhood [25]. Only one study, that of Chen *et al*. (1997) [39], found no significant association between the evolution of femoral neck BMD and the evolution of lean mass. The set of correlation coefficients is available in Figure 3, demonstrating a pooled correlation coefficient r of 0.33 (95% CI 0.16- 0.49), a value that is moderate but significant.

**Figure 3.**
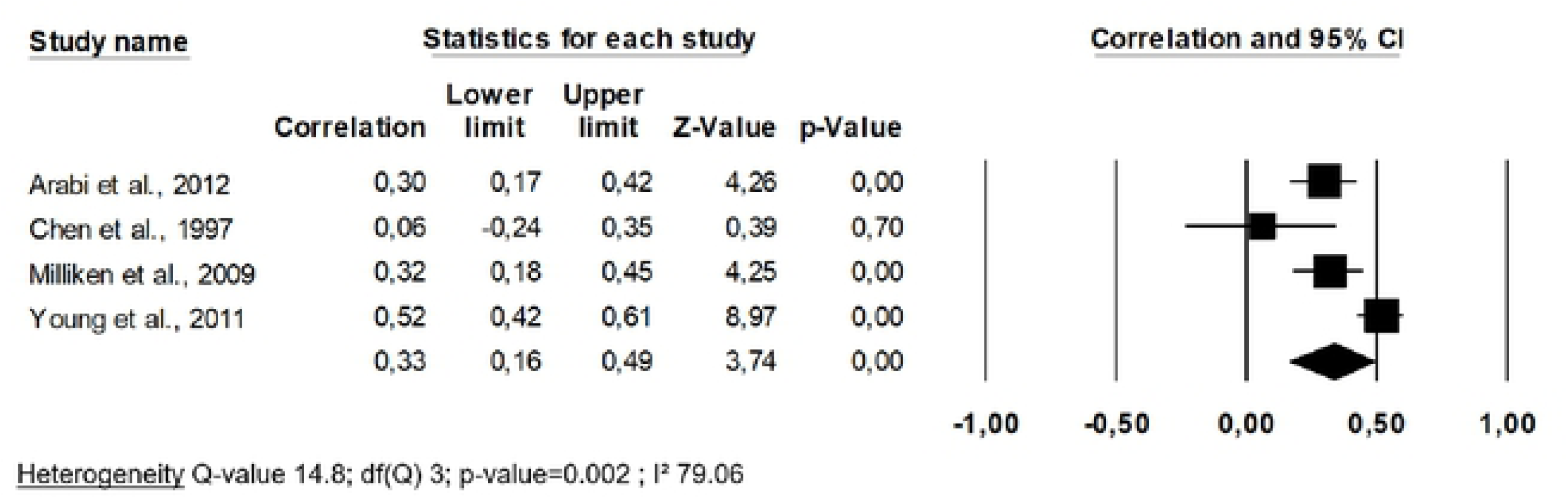
Association between changes in femoral neck bone mineral density and changes in lean mass: a meta-analysis

A subgroup analysis was not feasible since there was only one study in the ‘children’ group. Heterogeneity was found in this analysis (I^2^ = 79.06, p-value = 0.002). When we inserted moderators into the meta-regression, we observed that the quality level of the study, the mean age, the sex and the duration of follow-up did not have a significant impact on the pooled effect size (all p-values> 0.05). The Eggers’ regression test showed there was no publication bias (p-value = 0.24). A one-way analysis allowed us to determine that the study of Young *et al*. (2011) [25] had a significant influence on the meta-analysis model. Indeed, when this study was removed from the pooling, the heterogeneity was much lower and not significant (I^2^ = 22.36, p-value = 0.28). In this case, the pooled correlation coefficient raised up to 0.42.

### Lumbar Spine Bone Mineral Density and Lean mass

The four studies examining the prospective link between the evolution of lumbar spine BMD and the evolution of the lean mass were the same as the four studies described in the paragraph above, and they therefore have the same characteristics [25,29,36,39]. One study showed no significant association between the two entities tested [39]. The overall correlation coefficient r raised to 0.32 (95% CI 0.19-0.43), little heterogeneity between studies was observed (I^2^ = 60.91, p-value = 0.05) (Figure 4), and the one-way sensitivity analysis confirming this finding by demonstrating that the value of I^2^ remained in the same range.

**Figure 4.**
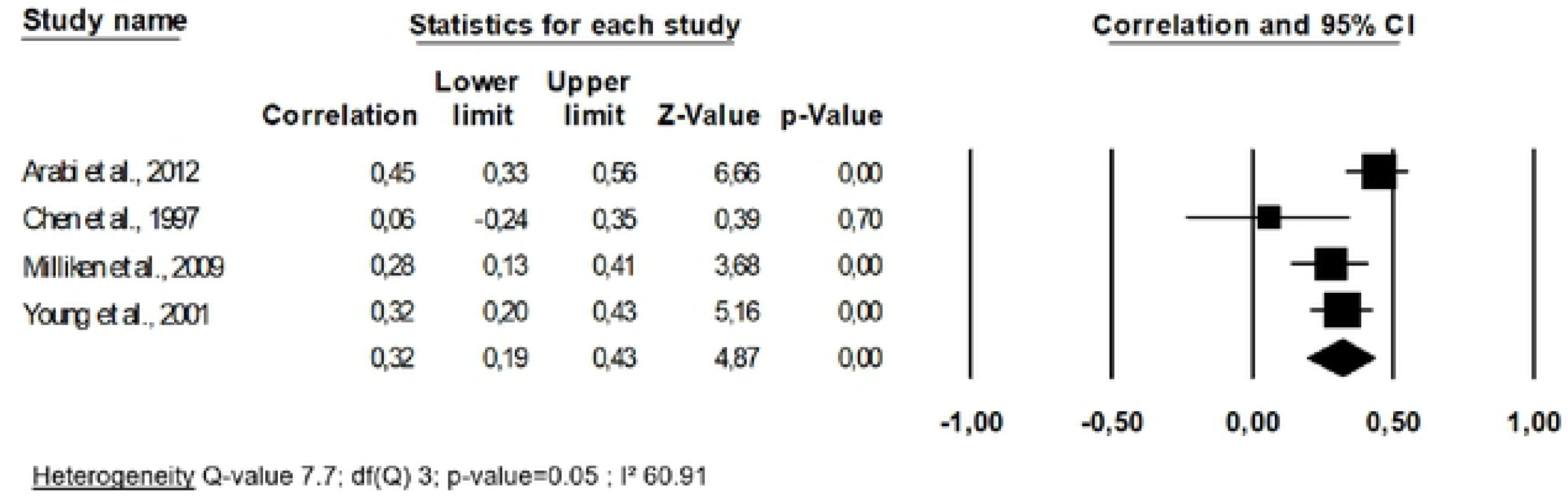
Association between changes in lumbar spine bone mineral density and changes in lean mass: a meta-analysis

When considering the quality level of the study, neither the mean age nor the duration of follow-up as moderators within a meta-regression, neither of those nor significantly stood out as moderators within the meta-regression (all p-values > 0.05). After applying Egger’s regression analysis, no publication bias was established (p-value = 0.33).

### Total Body Bone Mineral Density and Lean mass

The three studies included a total of 246 patients in this meta-analysis. Two studies were focused on childhood [31,34], and one in older subjects [39], and all were of good quality (NOS score = 8 points for all). One unique study by Chen *et al*. (1997) [39] showed no significant association between the changes in the two entities tested. When we pooled results of all studies, a correlation coefficient r of 0.53 was yielded (95% CI -0.23-0.89) and was not significant (p-value=0.16) (Figure 5). A large heterogeneity was found (p-value < 0.001). After performing the one-way analysis (i.e., one-to-one model withdrawal), heterogeneity remained high and significant.

**Figure 5.**
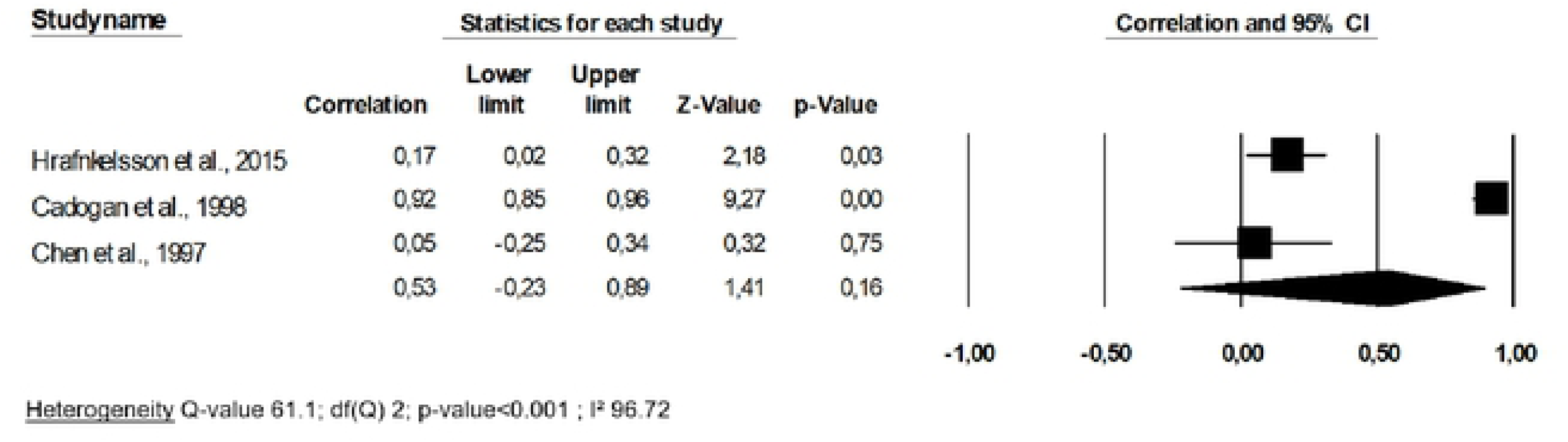
Association between changes in total body bone mineral density and changes in lean mass: a meta-analysis

A meta-regression could not be applied since too few studies were included. Finally, we did not find the presence of a publication bias, given that the p-value of Egger’s regression equaled 0.57.

### Bone Mineral Content and Lean mass

The study of Heidemann *et al*. (2015) [32] was not focused on the changes of BMD but rather in the changes of BMC. Therefore, this study required a separate analysis. The research was focused on children over a two-year follow-up period and appeared to be of fairly good quality (6 points out of 9). Heidemann *et al.* (2015) pointed out in their regression model that bone content accretion was significantly predicted by changes in lean mass (p-value <0.001).

### Relationship between Changes in Bone Mass and Changes in Muscle Strength

Three studies examined the link between the evolution of the BMD and that of muscular strength, in peri- or postmenopausal women [11,28,37]. However, since these three studies were performed by the same author, we did not perform a meta-analysis. In addition, two of these three studies appeared to focus on the same cohort of subjects. All three studies conclude that there is a statistically significant association between decreased muscle strength in postmenopausal women and bone loss (p-values <0.05). For these studies, no correlation coefficient was calculated, and the authors demonstrated an association simply based on differences in bone loss rate between those with increased grip strength and those with having decreased grip strength. These three studies are of good quality (7 out of 9 points).

Then, the study of Wang *et al*. (2007) [26], which was conducted on children two years of age, also showed that the change of BMC (but not the BMD) was significantly correlated with the change of volumetric contraction strength in the limbs (r ≥ 0.22, p-value < 0.05). This study was of quite good quality (6 out of 9 points).

### Relationship between Changes in Bone Mass and Changes in Physical Performance

Finally, the study by Kwon *et al.* (2007) [30], investigated the link between BMD changes and changes in physical performance, as measured by the 5-meter usual walking speed; this study was on older women and was methodologically excellent (9 points). This study concluded that a change in BMD was significantly related to a change in usual walking speed (correlation coefficient r = 0.21, p-value = 0.004). Indeed, a decline in BMD was significantly correlated to a decline in walking speed.

## Discussion

In this work we have synthesized data from the literature concerning the link between the evolution of skeletal status (specifically, BMD) and of various muscle components. No fewer than fifteen studies were included in our analysis, 10 of which were related to lean mass, 4 to muscle strength, and 1 to physical performance.

### Association between Changes in Bone Mineral Density and Changes in Lean mass

Using a meta-analysis approach, our results showed a significant association between the evolution of the bone mass and that of the mass muscle. The overall correlation coefficient r was significant and equal to 0.37 when considering bone hip density, 0.33 for femoral neck BMD and 0.32 for the lumbar spine. One notable exception, however, was that the link was not significant when measuring the bone density of the whole body (p-value = 0.16). However, this BMD measurement site does not appear optimal and would tend to overestimate the BMD and thereby underestimate osteopenia [40]. If one omits the unreliable results concerning the total body BMD, the magnitude of the correlation at the different sites seems to be the same (r varies from 0.32 to 0.37); this outcome suggests the same general trend of parallel evolution in the two entities of bone and muscle regardless of the measurement site.

A clear significant link was found (p-value < 0.001) between the two bone and lean mass, but it was moderate and thus one only partially explained the evolution of the other. Other factors should therefore be taken into account when considering the evolution of the masses of the bone-muscle unit (e.g., genetics; the genes that have been identified for their implications in sarcopenia [41] appear to be different from those involved in osteoporosis [42]). Evaluating the bone mass by using BMD or BMC measurements did not appear to influence the conclusions about the association between the two parameters.

The quite obvious relationship in development during the evolution of the bone mass and the lean mass could be explained mainly by mechanics: the muscular mass, which develops following a physical activity or an adapted dietary intake, will exert a net tension at the level of muscle insertion on the bone, resulting in a bone accruement [30,43]. During the maturing phase and the aging process, the opposite was observed: the physiological changes are linked to decreased physical activity, and a change in nutritional status leads to a decline in lean mass, thus no longer exerting its mechanistic effect on the bones of the body and causing a simultaneous decline in bone mass. Our study shows that this link is more marked during childhood and therefore during the developmental process (r = 0.43) than during the aging process (r = 0.34). We hypothesize that this significant difference in effect (p <0.001) may be related to multiple other influencing factors (such as diet, gene expression, peak bone and lean mass acquired during childhood, …) that have had the time to impact musculoskeletal health during adulthood and aging.

### Association between Changes in Bone Mineral Density and Changes in Muscle Strength

Few studies have been identified regarding this aspect: 3 studies by the same author, and one in children. In both situations, we found significant associations. Although no meta-analysis could be done, these studies revealed a link between the evolution of muscle strength and that of the BMD. Concluding with an association seems coherent if we emit the assumption of mechanical strain on bone tissue metabolism [12,44].

### Association between Changes Bone Mineral Density and Changes in Muscle Performance

We have only one study regarding the relationship of the evolution of the BMD compared to that of physical performance, although this issue deserves further investigation. There are many studies on whether physical performance (weak or good) is a determinant of low or good bone mass [45] but not on if there is a parallel evolution between the two entities, although this is a more preventive idea. However, in the only study that highlighted the link between bone mass and physical performance, the correlation was significant albeit moderate (r = 0.21). This significant association could be explained by the direct link between the physical function (and by this, the performance) and the muscular action on the bone, as emphasized previously by several authors [30,43].

### Strengths and Limitations

This study is the first to employ a meta-analysis to synthetize the results from studies examining an association between the parallel evolution of bone and muscle status. One asset of our study is that we looked in multiple databases, although some databases could not be consulted following logistical barriers. All processes have been rigorous, using the PRISMA statements to ensure a good level of reporting of our research. Our present analysis, however, has certain limitations, so it is necessary to consider its interpretation in the context of these limitations. First, even though we made every effort to contact the authors to obtain the missing data that was essential for the quantitative synthesis of the results, the majority of the authors did not answer our calls. Therefore, we had to resort to a technical imputation of the correlation coefficient r from the coefficient β. Although the technique we used is well-funded, it does not provide real data and can therefore lead to heterogeneity. However, this technique gives us precise data that allow us to pool the results rather than omit these studies from the research; such an omission would be damaging. Then, for the obvious reasons of statistical pooling of our results, we used a correlation coefficient, a measure that does not consider the influence of potential confounding factors.

We also identified the potential sources of heterogeneity among the included references: the sex of the subjects studied, the quality of the studies, and the diverse duration of follow-up and age-group of interest (i.e., differences in protocols). However, we have taken these parameters into account by using a random-effect model and performing subgroup or meta-regression analyses, showing no effect of duration of follow-up but significant differences in outcomes between children and adults (i.e., the association is significantly greater in children than in adults). We hypothesize that this heterogeneity could also be induced by the small number of studies that were included in each of the meta-analyses, and other good quality studies should be conducted to reinforce our analysis, despite the fact that this statement seems clear (i.e., although there is indeed a significant association, it seems moderate). It should also be noted that there remains a possibility that our analysis has necessarily excluded unpublished studies concerning our theme, representing a threat to its validity. However, the publication bias was evaluated and did not show any bias at this level even if some relevant papers could always been missed. Finally, it should also be recognized that the instrument that was used to measure the methodological quality of the studies also had its limitations, particularly in terms of its interpretation. This could influence our descriptive but also statistical analysis (i.e., the results of the meta-regression).

## Conclusion

Our analysis definitively highlighted a net relationship between the changes, both during childhood and during the aging process, of bone mass and of markers of muscle function. Thus, it is possible that the development and the decline of those two entities deserve to be considered as one unique musculoskeletal entity. On this basis, health-enhancing, preventive and therapeutic interventions could be developed by targeting these two structures at the same time.

## Acknowledgments

We warmly thank Dr. Macdonald and Dr. Liu-Ambrose for providing us with important information regarding their articles, allowing us to conduct our meta-analysis in the best and most accurate way possible. Additionally, there is no funding to declare.

## Authors’ roles

Study design: ML, CB, OB, and ND. Study conduct: ML, and CB. Data collection: ML, and CB. Data analysis: ML. Data interpretation: ML, CB, JYR, OB. Drafting manuscript: ML. Revising manuscript content: ML, CB, ND, JYR, and OB. Approving final version of manuscript: ML, CB, ND, JYR, and OB. ML takes responsibility for the integrity of the data analysis.

## Notes

**Disclosures** All authors declare no conflict of interest

